# Unique ecology of co-occurring functionally and phylogenetically undescribed species in the infant oral microbiome

**DOI:** 10.1101/2025.06.03.657575

**Authors:** Nicholas Pucci, Amke Marije Kaan, Joanne Ujčič-Voortman, Arnoud P. Verhoeff, Egija Zaura, Daniel R. Mende

**Affiliations:** Department of Medical Microbiology & Infectious Diseases, Amsterdam UMC, Location AMC at University of Amsterdam, Amsterdam, The Netherlands; Department of Preventive Dentistry, Academic Center for Dentistry Amsterdam, University of Amsterdam and Vrije Universiteit Amsterdam, Amsterdam, The Netherlands; Sarphati Amsterdam, Public Health Service Amsterdam, Amsterdam, The Netherlands; Department of Sociology, University of Amsterdam, Amsterdam, The Netherlands; Human Biology-Microbiome-Quantum Research Center (WPI-Bio2Q), Keio University, Tokyo, Japan

## Abstract

Early-life oral microbiome development is a complex community assembly process that influences long-term health outcomes. Nevertheless, microbial functions and interactions driving these ecological processes remain poorly understood. In this study, we analyze oral microbiomes from a longitudinal cohort of 24 mother-infant dyads at 1 and 6 months postpartum using shotgun metagenomics. We identify two previously undescribed *Streptococcus* and *Rothia* species to be among the most prevalent, abundant and strongly co-occurring members of the oral microbiome of six-month-old infants. To explore the underlying genomic traits enabling this unique lifestyle, we leveraged metagenome-assembled genomes (MAGs) and genome-scale metabolic models (GEMS). Comparative analyses revealed specific genomic and functional characteristics relative to other closely related species and highlighted unique functional features, including genes encoding adhesins and carbohydrate-active enzymes (CAZymes). The observed co-occurrence patterns were further supported by predicted metabolic interactions within a network of co-occurring oral taxa. Metabolic modeling identified potential exchange of key nutrients, particularly malate and lysine, between these species, suggesting metabolic cross-feeding interactions that may explain their co-occurrence across infant oral microbiomes. Overall, this study provides key insights into the functional adaptations and microbial interactions shaping early colonization in the oral cavity.

**Author Summary:** We investigated how infant oral bacterial communities develop during their first six months of life, with the aim to understand which microbes colonize, how they establish themselves and why they succeed together. Using high throughput DNA sequencing techniques, we analyzed oral samples from 24 mother-infant pairs at one and six months after birth.

We found two abundant, but previously unknown bacterial species (one Streptococcus and one Rothia) at six months of age. These bacteria consistently appear together across different babies, suggesting they may depend on each other for survival and growth.

By reconstructing the genomes of these bacteria directly from our samples, we discovered specific genetic features that help explain their success in the infant mouth. Streptococcus carries genes that allow it to break down nutrients from breast milk. Rothia has genes that help it rapidly build cell membranes and protect against harmful molecules, while producing nutrients that Streptococcus needs. We predict these bacteria exchange key nutrients like malate and lysine, creating a mutually beneficial partnership. The bacteria seem to cooperate and are predicted to exchange key nutrients like malate and lysine which may help maintain a healthy oral environment by regulating acidity levels, potentially protecting against tooth decay.

## Introduction

The first bacteria (‘animalcules’) Antonie van Leeuwenhoek observed in the 17th century were oral microbes. In centuries afterwards, oral microbes were found to be involved in a range of conditions, such as dental caries, periodontal diseases and oral candidosis (1). An understanding of the beneficial role that the microbes constituting the oral microbiome play was realized much later. A balanced oral microbiome promotes oral health by serving as a protective barrier, preventing infections and colonization of oral surfaces by harmful pathogens as well as regulating oral pH (2,3). Beyond oral health, the oral microbiome has significant implications for systemic health (4), contributing to vasodilation and blood pressure regulation through nitrogen and enterosalivary nitrate metabolism (5). Though its importance is undeniable, our knowledge about the oral microbiome, particularly its development during infancy, remains limited, and public metagenomics datasets remain scarce.

Most of the existing knowledge about the infant oral microbiome stems from 16S rRNA gene amplicon surveys, which revealed the genus composition of the oral microbiome development through a predictable succession of microbial colonization, where early colonizers such as *Streptococcus* and *Actinomyces* adhere to oral tissues thanks to their adhesin repertoire (6–11). These pioneers subsequently form a foundation that supports the establishment of secondary colonizers, including *Veillonella* and *Rothia* (12). Developmental milestones, such as primary dentition, and environmental exposures are often accompanied by further compositional shifts (8). While these coarse grained taxonomic succession patterns have been described, the functional potential and interactions within these microbial communities, particularly during the formative months of life, remain poorly characterized. Yet, such knowledge about the developing oral microbiome would be crucial given the potential persisting effects on oral and systemic health later in life (8,13,14).

Microbial interactions fundamentally shape microbiome structure and function (12,15), yet little is known about their contribution to the stability and resilience of the developing oral microbiome. Typically, these interactions are a combination of spatial organization (16,17), complex metabolic dependencies and cross-feeding mechanisms (18) leading to mutualistic, competitive or commensal relationships (3,19–21), while driving community-level processes such as pH regulation (22) and nitrate reduction (23).

Understanding the genomic basis of these synergistic activities, including metabolic pathways and genetic determinants, is crucial for deciphering how microbial communities establish and maintain functional stability during infant oral microbiome development.

Methodological advances allowing for the reconstruction of genomes directly from metagenomics, and their interrogation through comparative approaches (metapangenomics) (24) have the potential to provide insight into the functional potential and metabolic interactions of the developing microbiome. For example, genome-resolved metagenomic approaches allowed Utter et al. (24) to have strain-level resolution and identification of accessory genes conferring adaptive advantages to *Rothia* and *Haemophilus* spp. to specific oral niches in healthy adults. Further, novel tools allow for the generation of accurate metabolic models directly from genomes (25,26), which allows the prediction of substrate utilization and metabolite production. Community models combining the metabolic models of multiple co-occurring microbes can provide further insight into metabolic interactions and cross-feeding, therefore elucidating the metabolic basis for the synergistic activities and successional patterns observed during oral microbiome development (27–29).

Here, we performed shotgun metagenomics on a time series of 24 infants and their mother’s oral microbiome and identified novel, co-occurring species using a meta-pangenomics approach. Analysis of a time-series of tongue dorsum and dental biofilm samples collected from children up to six months of age and their mothers as part of the prospective Amsterdam Infant Microbiome Study (AIMS) revealed previously undescribed *Streptococcus* and *Rothia* species to be predominant members of the infant oral cavity. By leveraging metagenome-assembled genomes (MAGs), we characterize genomic features and functional attributes of these species to explain their adaptations to the infant oral cavity. In order to interpret their co-occurrence patterns we further decipher their metabolic interactions within an oral species network. Overall, our findings highlight important microbial ecological dynamics in the developing oral ecosystem of infants and establish a foundation for understanding how early microbial interactions help shape the infant oral microbiome.

## Results

### Undescribed *Streptococcus* and *Rothia* species emerge as predominant in six-month infant oral microbiome

We investigated the oral microbiome of 24 mother-infant pairs from the AIMS cohort, focusing on full-term, vaginally-born infants with no reported antibiotic exposure (Table S1). By six months, 87% of AIMS infants (21 out of 24) had obtained their first teeth, allowing collection of dental biofilm. Shotgun metagenomics sequencing (mean depth tongue biofilm: 20.5M reads per sample; mean depth dental biofilm: 15.7M reads per sample; Figure S1) was performed on samples collected from tongue dorsum swabs from mothers (successfully sequenced samples: n=22) and infants both 1 (n=9) and 6 months (n=18), and dental biofilm from mothers (n=21) and infants at 6 months (n=14). All analyzed one-month-old infants were exclusively breastfed. By six months, we observed diverse feeding patterns including breast- (n=8), mixed-(n=7) and formula feeding (n=5), with 95% (21 of 22) of infants being introduced to solid foods (Table S1).

Metagenomic profiling revealed the *Streptococcus* genus to be the most abundant and prevalent taxonomic group in the oral microbiome of AIMS infants at both 1 and 6 months of age. However, we observed a notable species-level compositional shift over time, with previously undescribed species emerging by six months. At one month, infant tongue dorsum hosted an average of 19 (±6) bacterial species (Fig. 1B) and was predominantly composed of *Streptococcus* (relative abundance±standard deviation, SD 57.0±6.8%), *Veillonella* (9.0±2.0%), *Lactobacillus* (6.1±3.3%) and *Rothia* (4.9±1.8%) species (Fig. 1A). By six months, we observed an increase in bacterial species richness (tongue dorsum 43±17, dental biofilm 43±14) with an overall shift in tongue microbiome composition relative to one month (Fig. 1A-C). Notably, previously undescribed *Rothia* (GTDB: sp902373285: relative abundance=9.2±2.2%, prevalence=61%) and *Streptococcus* (GTDB: sp000187445: 5.6±1.9%, 66%) species, hereon referred to as *Streptococcus* AIMSoral1 and *Rothia* AIMSoral2, emerged as abundant and prevalent bacteria in the oral cavity, particularly in the tongue biofilm (Fig. 1B). Dental biofilm composition at six months (despite the emergence of teeth as a new niche) largely resembled the tongue biofilm composition (Fig. 1C). Milk feeding type did not significantly influence community composition (Fig. S3). Maternal dental and tongue biofilm communities were markedly different (Fig. 1C) and strain sharing was found to be rare with only two events found across the whole cohort (Table S2), both between mothers and their respective one-month-old infants.

**Figure 1.**
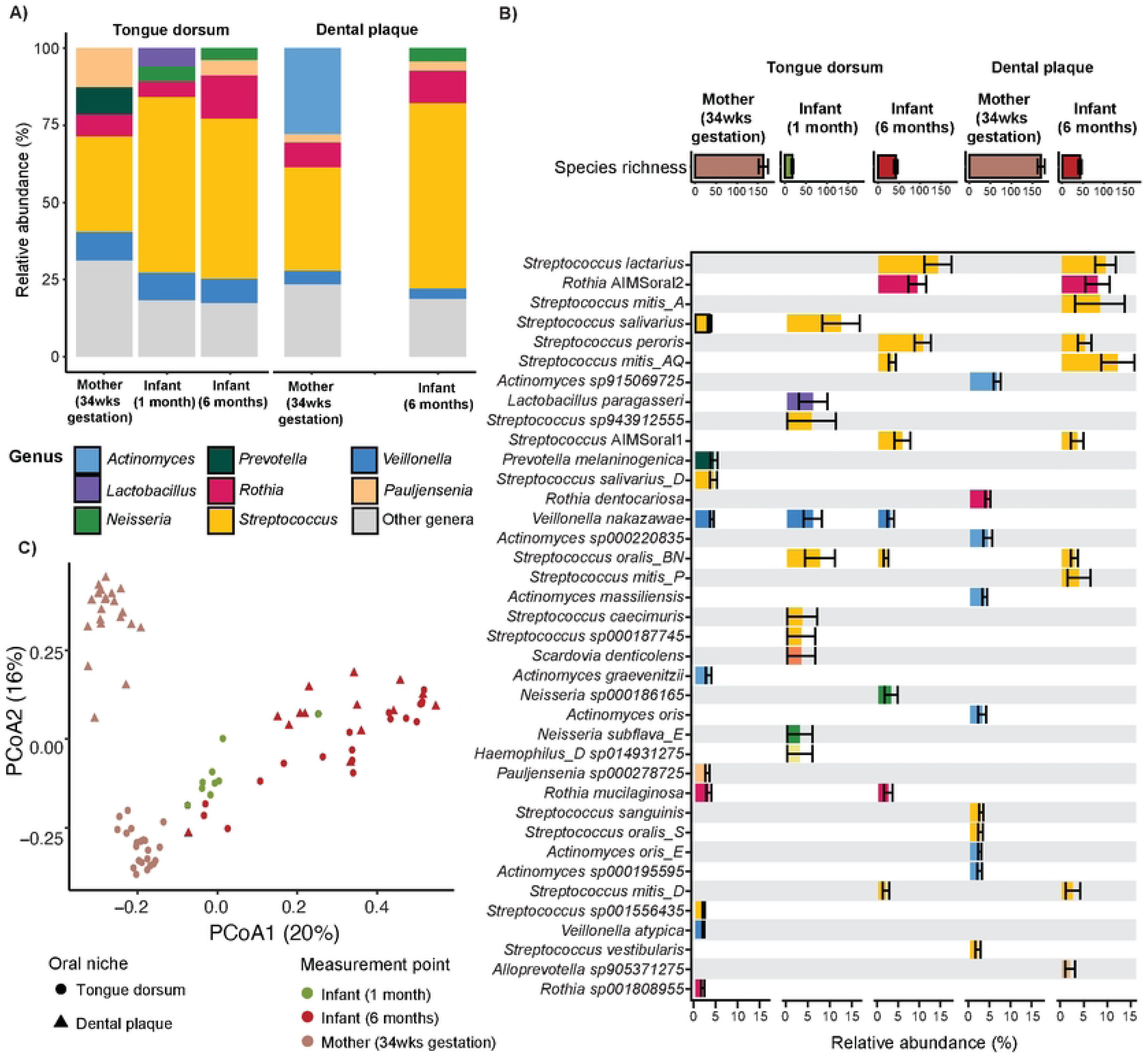
Oral microbiome abundance and composition of AIMS mothers and their infants. **A)** Relative abundance of the 5 most abundant bacterial genera detected in the tongue dorsum and dental plaque of mothers (34wks gestation) and their infants (one and six months of age). **B)** Relative abundance (mean±SD) of the 15 most abundant bacterial species detected in the tongue dorsum and dental plaque of mothers (34wks gestation) and their infants (one and six months of age). Species displayed include all taxa that ranked among the 15 most abundant in at least one sample group or time point. Bars are coloured by genus. **C)** Principal coordinate analysis (PCoA) of tongue dorsum (circles) and dental plaque (triangles) microbiomes from AIMS mothers (34wks gestation=brown) and their infants (one month=green; six months=red).

### *Streptococcus* AIMSoral1 and *Rothia* AIMSoral2 show significant co-occurrence in infant oral ecosystems

To elucidate potential species interactions, we inferred pairwise co-occurrence patterns across ages by combining hierarchical clustering and Spearman’s rank correlations for tongue and dental biofilm samples, separately. As the most abundant species in the infant oral microbiomes were almost all absent from the maternal oral microbiomes (Fig. 1B), we present co-occurrence network results for infant samples only. In one-month-old edentate infants, we detected 31 species pairs with significant correlations involving 28 (of 32) abundant (relative abundance≥1%) tongue biofilm species (Fig. S4A, Table S3). Among these correlations, we found few involving predominant tongue species (relative abundance≥5%, prevalence>30%). *L. paragasseri* showed a significant pairwise correlation with *R. mucilaginosa_A,* only (Spearman’s rank correlation: R=0.87, *p*=0.002). Similarly, *S. oralis*_BN showed a significant, but negative correlation with *V. nakazawae* (Spearman’s rank correlation: R=-0.88, *p*=0.002), while *S. salivarius* showed no significant correlations.

By six months of age, we detected 24 and 21 significant pairwise abundance correlations involving 14 (out of 17) and 17 (out of 24) species across tongue and dental biofilms, respectively. *Rothia* AIMSoral2 and *Streptococcus* AIMSoral1 displayed the strongest positive co-occurrence among abundant bacteria in the infant tongue biofilm (Spearman’s rank correlation: R=0.68, p=0.00008; Fig. S4B, Table S3). *Rothia* AIMSoral2 and *Streptococcus* AIMSoral1 significantly co-occurred with nine and four other species, respectively (Fig. S4B). Notably, both species exhibited significant abundance correlations with undescribed *Veillonella* (GTDB: sp018367495) and *Pauljensenia* (GTDB: sp900541895) species on the tongue dorsum, hereon referred to as *Veillonella* AIMSoral3 and *Pauljensenia* AIMSoral4. These species also correlated significantly with each other, forming a co-occurrence network of four species. Neither of these species was detected in maternal or one month oral samples.

In dental biofilm, the significant co-occurrence between *Rothia* AIMSoral2 and *Streptococcus* AIMSoral1 persisted, but was weaker (R=0.40, p=0.04). Both species also showed strong correlations with *Veillonella* AIMSoral3 (*Rothia* AIMSoral2: R=0.54, p=0.004; *Streptococcus* AIMSoral1: R=0.54, p=0.01) (Fig. S4C, Table S3). Overall, abundance of these species was not significantly influenced by feeding type (Fig. S4).

### Genomic relatedness analyses reveal undescribed *Streptococcus* and *Rothia* to be novel species

Among identified species in the developing oral microbiome, the undescribed *Streptococcus* AIMSoral1 and *Rothia* AIMSoral2 were selected for functional characterization due to their high abundance (≥5%) and prevalence (>60%) across infant oral cavities. To confirm metagenomic taxonomic profiling of these species, we reconstructed metagenome-assembled genomes (MAGs).

Metagenomic binning from maternal and infant oral samples yielded 20 *Streptococcus* (completeness, mean±SD=64.15±12.17%; contamination, mean±SD=2.11±2.84%) and 51 *Rothia* (91.24±13.32%; 0.91±1.27%) medium and high-quality MAGs (>50% completeness and <10% contamination), of which five were taxonomically assigned to *Streptococcus* AIMSoral1 and 15 to *Rothia* AIMSoral2 using GTDB-tk (Table S5). To verify these taxonomic assignments, we reconstructed phylogenomic trees based on single-copy core gene protein sequences (Fig. 2), incorporating reference genomes from public databases (Table S5).

**Figure 2.**
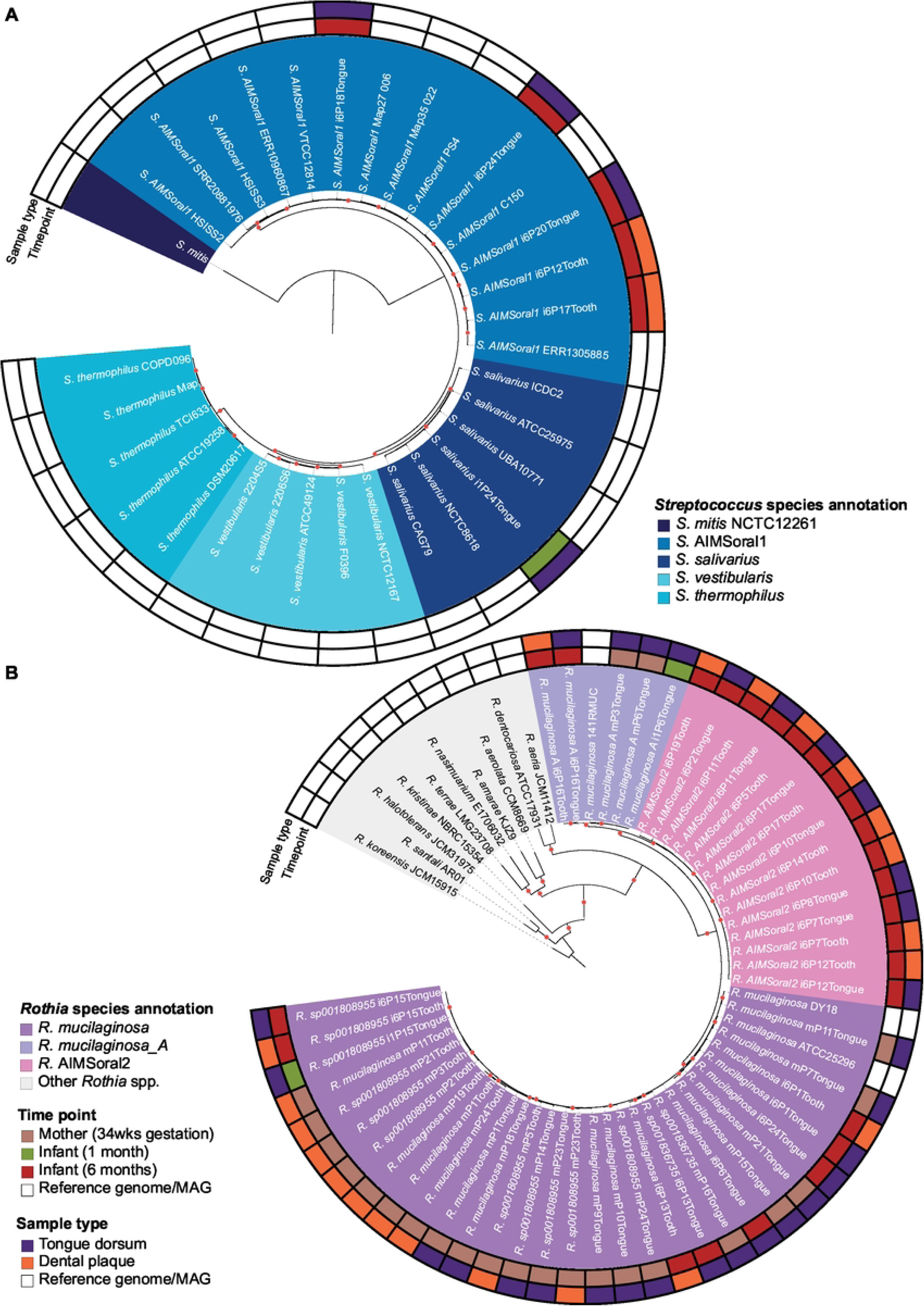
Phylogenomic trees of *Streptococcus salivarius* clade and *Rothia* genus, highlighting the undescribed species *Streptococcus* AIMSoral1 and *Rothia* AIMSoral2. **A)** Phylogenomic tree of the *salivarius* clade of the *Streptococcus* genus containing 33 genomes (reference genomes=27, MAGs=6). **B)** Phylogenomic tree of the *Rothia* genus containing 63 genomes (reference genomes=13, MAGs=50). Genome names are colored by species identity as defined by genetic relatedness analyses. Rings indicate the time point (mother 34wks gestation, infant one and six months) and sampling location (tongue dorsum and dental plaque). Red circles indicate bootstrap values≥0.9.

Phylogenomic analysis of 89 *Streptococcus* species (110 genomes) placed *Streptococcus* AIMSoral1 within the *salivarius* group with high confidence (Fig. S5). To refine this placement, we employed a pangenomic approach focusing specifically on the *salivarius* group, constructing a phylogenomic tree from single-copy core genes (Fig. 2A). This targeted analysis included 27 reference genomes and six MAGs: *S. salivarius* (n=6), *S. vestibularis* (n=5), *S. thermophilus* (n=5) and *Streptococcus* AIMSoral1 (n=15). The phylogenetic analysis revealed *Streptococcus* AIMSoral1 to form a distinct, well-supported monophyletic clade, representing a sister clade to the *salivarius* group. This observation was supported by average nucleotide identity (ANI, species threshold=95%) (32) analyses, confirming that *Streptococcus* AIMSoral1 represents a species-level entity, with 97.3% to 98.6% ANI within its cluster, but lower ANI to phylogenetically neighboring species (*S. salivarius*: 88.6-89.3%, *S. vestibularis*: 88.3-88.5%, *S. thermophilus*: 87.1-87.4%; Fig. S6A).

The *Rothia* phylogenomic tree included 63 genomes (reference genomes=13, MAGs=50) and placed the undescribed *Rothia* AIMSoral2 in a monophyletic clade, closely related to *R. mucilaginosa*_A and *R. mucilaginosa* (33) (Fig. 2B).

For *Rothia*, ANI analyses also supported *Rothia* AIMSoral2 as a species-level entity, showing 97.0% to 99.9% ANI within its cluster, but markedly lower ANI with phylogenetically neighboring *R. mucilaginosa*_A (90.2-91.4%) and *R. mucilaginosa* (86.9-88.5% ANI) (Fig. S6B).

### Adhesion and carbohydrate utilization differentiate *Streptococcus* AIMSoral1 from other infant-associated species

To uncover functional traits driving the abundance and persistence of *Streptococcus* AIMSoral1 in the infant oral cavity, we compared its pangenome, metabolic capacity, and adhesion potential to other infant-associated *Streptococcus* species (see Methods: Public reference genomes acquisition).

The *Streptococcus* pangenome included 57 genomes (Table S5) from six infant-associated species: *Streptococcus* AIMSoral1 (n=14), *S. lactarius* (n=12), *S. salivarius* (n=6), *S. oralis* (n=9), *S. mitis* (n=12) and *S. peroris* (n=3). *Streptococcus* AIMSoral1 genomes averaged 1.9±0.9Mb (average±SD), carrying roughly 1930±98 genes. Gene-cluster (GC) analysis showed largely species-driven clustering, with *Streptococcus* AIMSoral1 displaying distinct, but close genomic similarity to *S. salivarius* (Fig. S7). The pangenome was further analyzed to reveal metabolic capabilities of *Streptococcus* AIMSoral1 supporting its predominance during infant weaning.

Adhesion potential, assessed via lectin-encoding gene copy number, largely mirrored species identity (Fig. 3). The adhesin repertoire of *Streptococcus* AIMSoral1 showed high similarity to *S. salivarius* and *S. lactarius*, including enrichment in *fimA* lipoprotein genes (present in *Streptococcus* AIMSoral1 and *S. lactarius*, but absent in other species; Fig. 3, Table S10).

**Figure 3.**
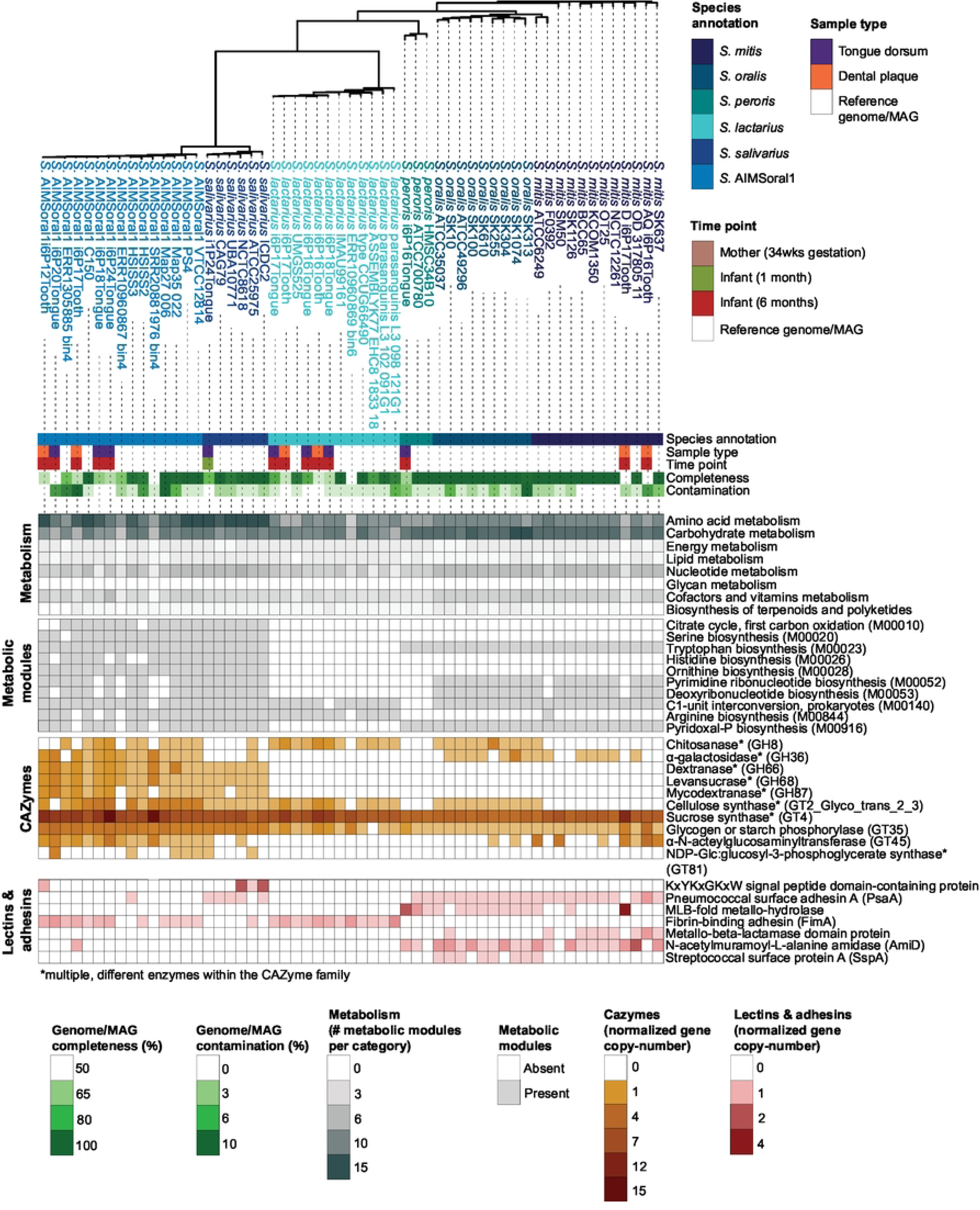
Overview comparative genomics analyses outputs for *Streptococcus* AIMSoral1 and infant-associated *Streptococcus* species. Phylogenomics tree based on single-copy core genes of 57 infant-associated *Streptococcus* genomes, color-coded by species annotation: *Streptococcus* AIMSoral1, *S. salivarius*, *S. lactarius*, *S. peroris*, *S. oralis* and *S. mitis*. Genome completeness and contamination metrics are displayed for each genome, together with sampling location and time point of origin. Heatmaps below the tree show for each genome comparative genomics analyses results, highlighting metabolic modules/genes significantly enriched in *Streptococcus* AIMSoral1 relative to other infant-associated *Streptococcus* species. From top to bottom, panels display: metabolic module abundance by broad metabolic pathway (Metabolism=gray palette), metabolic modules (light gray), CAZyme- (orange palette) and lectin/adhesin- (red palette) encoding genes enriched in *Streptococcus* AIMSoral1 relative to other infant-associated *Streptococcus* spp.

Carbohydrate-active enzymes (CAZymes) analysis revealed *Streptococcus* AIMSoral1 harbors significantly higher copy numbers of eight glycosyltransferases (GTs), including α-N-acetylglucosaminyltransferase (GT45) and starch phosphorylases (GT35), as well as four glycoside hydrolases (GHs), such as GH66, GH68, and GH87 families, which were shared between *Streptococcus* AIMSoral1 and *S. salivarius* (Fig. 3, Table S10). Additionally, *Streptococcus* AIMSoral1 had significantly more GH8, GH36, and GT2_Glyco_trans_2_3 genes compared to *S. salivarius*, suggesting distinct carbohydrate-utilization strategies. Overall, *Streptococcus* AIMSoral1 and *S. salivarius* have fewer genes involved in carbohydrate metabolism than other infant-associated species (Fig. 3).

Metabolic enrichment analysis showed *Streptococcus* AIMSoral1 and *S. salivarius* to be significantly enriched in genes for amino acid metabolism compared to other infant-associated species (Fig. 3). Notably, only *Streptococcus* AIMSoral1 and *S. salivarius* encoded the full arginine biosynthesis pathway from glutamate (M00028, M00844; Table S6), a downstream metabolite of the citrate cycle (also enriched; M00010; Table S6) and major component of breast/formula milk. No metabolic module was exclusively enriched in *Streptococcus* AIMSoral1.

### Carbohydrate utilization and fatty acid biosynthesis differentiate *Rothia* AIMSoral2 from other *R. mucilaginosa* species

We compared genomic traits of *Rothia* AIMSoral2 to infant-associated *Rothia* species to uncover metabolic adaptations supporting its predominance across 6-month-old infant oral cavities.

The *Rothia* pangenome included 53 MAGs and reference genomes, from three species (as annotated by GTDB-tk), retrieved from infants in our study: *R. mucilaginosa* (30 MAGs, 2 reference genomes), *R. mucilaginosa_A* (5 MAGs, 1 reference genome), and *Rothia* AIMSoral2 (15 MAGs, no reference genomes available). *Rothia* AIMSoral2 genomes averaged 2.1±0.1Mb and carried 1760 (±37) genes. In line with phylogenomic analysis, these genomes form a consistent clade based on gene presence/absence, showing greater similarity to *R. mucilaginosa*_A than other *R. mucilaginosa* strains (Fig. S8).

All infant-associated *Rothia* species carried genes-encoding for CBM50 domains (also known as LysM; Table S10), associated with binding bacterial peptidoglycans (34). No matches were found in the LectomExplore database for *Rothia* lectins, suggesting limited host-surface adhesion potential. Further CAZyme analysis showed *Rothia* AIMSoral2 to be enriched in genes encoding for, among others, GT2, GT4, and GT11, involved in EPS production, biofilm formation and surface adhesion (35–39) (Table S10). Additionally, *Rothia* AIMSoral2 carried significantly higher copy-numbers of redox-active enzyme-encoding genes (Auxillary Activities: AA1 and AA1_1/2; Fig. 4 and Table S10), co-acting with CAZymes.

**Figure 4.**
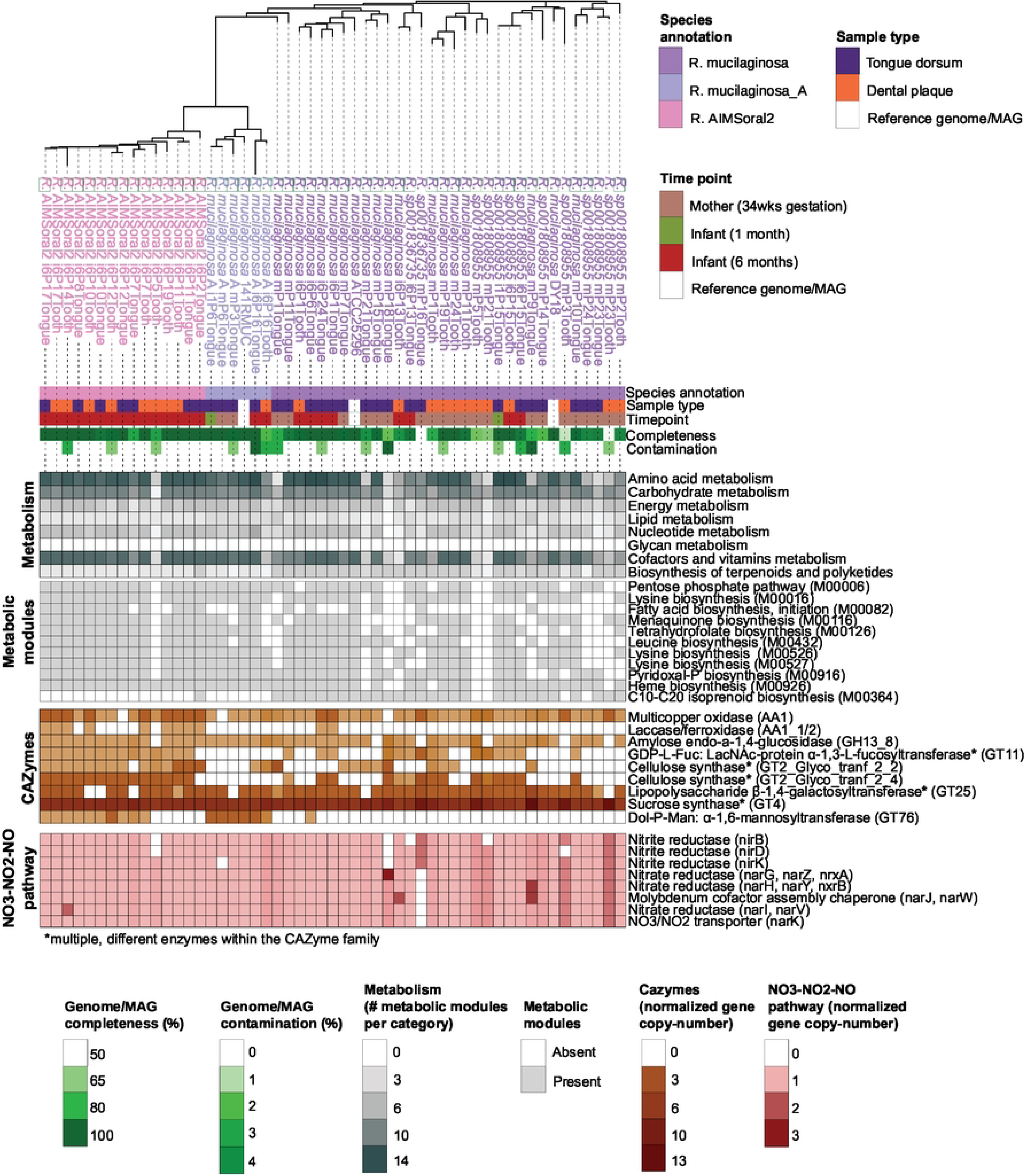
Overview comparative genomics analyses outputs for *Rothia* AIMSoral2 and infant-associated *Rothia* species. Phylogenomics tree based on single-copy core genes of 53 infant-associated *Rothia* genomes, color-coded by species annotation: *Rothia* AIMSoral2, *R. mucilaginosa* and *R. mucilaginosa_A*. Genome completeness and contamination metrics are displayed for each genome, together with sampling location and time point of origin. Heatmaps below the tree show for each genome comparative genomics analyses results, highlighting metabolic modules/genes significantly enriched in *Rothia* AIMSoral2 relative to other infant-associated *Rothia* species. From top to bottom, panels display: metabolic module abundance by broad metabolic pathway (gray palette), metabolic modules (light gray), CAZyme-encoding (orange palette) and enterosalivary pathway-associated (red palette) genes enriched in *Rothia* AIMSoral2 relative to other infant-associated *Rothia* spp.

We further investigated genes involved in the nitrate-nitrite-nitric oxide (NO_3_-NO_2_-NO) enterosalivary pathway in infant-associated *Rothia* species. All infant-associated *Rothia* species carried genes involved in the respiratory denitrification of NO_3_ to NO and, in part, intracellular ammonia (NH_3_) production. Further functional analyses revealed *Rothia* AIMSoral2 to carry significantly higher numbers of genes associated with cofactors and vitamin biosynthesis compared to *R. mucilaginosa,* but not *R. mucilaginosa_A* (Fig. 4). Notably, the key process of fatty acid biosynthesis initiation (M00082) was significantly enriched in *Rothia* AIMSoral2 (Fig. 4; Table S7).

### *Streptococcus* AIMSoral1 shows higher metabolic complementarity to *Rothia* AIMSoral2 than other *Streptococcus* species

To investigate whether the co-occurrence of *Streptococcus* AIMSoral1 and *Rothia* AIMSoral2 at six months of age corresponded with their metabolic interdependence, we reconstructed GEMS for infant-associated *Streptococcus* and *Rothia* species using the PhyloMInt tool (Methods: Metabolic complementarity/interaction potential).

Overall, *Streptococcus* species exhibited comparatively greater metabolic complementarity (MI_complementarity_) with *Rothia* than viceversa (Fig. S9A, Table S13), though we found significant differences among individual species. *Rothia* AIMSoral2 exhibited, comparatively, the highest MI_complementarity_ with *S. lactarius* and *Streptococcus* AIMSoral1. Despite *Streptococcus* AIMSoral1 and *S. salivarius* having similar metabolic profiles, *S. salivarius* showed the lowest MI_complementarity_ with all *Rothia* species. While *Rothia* AIMSoral2 and *R. mucilaginosa_A* had comparable MI_complementarity_ patterns, *R. mucilaginosa_A* showed the highest MI_complementarity_ with all *Streptococcus* species. Notably, *Streptococcus* AIMSoral1 (and *S. peroris*) exhibited significantly greater MI_complementarity_ with all *Rothia* species than other *Streptococcus* species.

Using the PhyloMInt PTM tool, we identified metabolites each species could produce and utilize, allowing us to assess potential differences in metabolic interactions between *Rothia* AIMSoral2 and *Streptococcus* AIMSoral1 compared to their interactions with other infant-associated species. We predicted multiple potential metabolic interactions between *Rothia* AIMSoral2 and *Streptococcus* AIMSoral1, but these were often shared with other species (Fig. S9B, Table S14).

### Genome-scale predictions of metabolic interactions in an oral bacterial network

We hypothesized that metabolic interactions facilitate the co-occurrence network comprising S*treptococcus* AIMSoral1, *Rothia* AIMSoral2, *Veillonella* AIMSoral3 and *Pauljensenia* AIMSoral4 (Fig. 5A). To test this hypothesis we predicted metabolic interactions among these species. Although we lacked sufficient *Veillonella* AIMSoral3 and *Pauljensenia* AIMSoral4 MAGs for comprehensive comparative genomics, metabolic prediction was still possible using available reference genomes. Using GEMS, we inferred MI_complementarity_ based on the highest quality available MAG or reference genome (*see* Methods: Prediction of metabolic complementarity and interaction potentials; Table S13). Comparatively, all species exhibited the lowest MI_complementarity_ to *Streptococcus* AIMSoral1, while the latter exhibited the highest MI_complementarity_ with *Pauljensenia* AIMSoral4 and *Rothia* AIMSoral2 (Fig. 5B). Both *Rothia* AIMSoral2 and *Pauljensenia* AIMSoral4 were comparatively highly complementary to *Veillonella* AIMSoral3. Notably, *Veillonella* AIMSoral3 showed the strongest MI_complementarity_ with *Rothia* AIMSoral2, suggesting a key metabolic relationship between these species.

**Figure 5.**
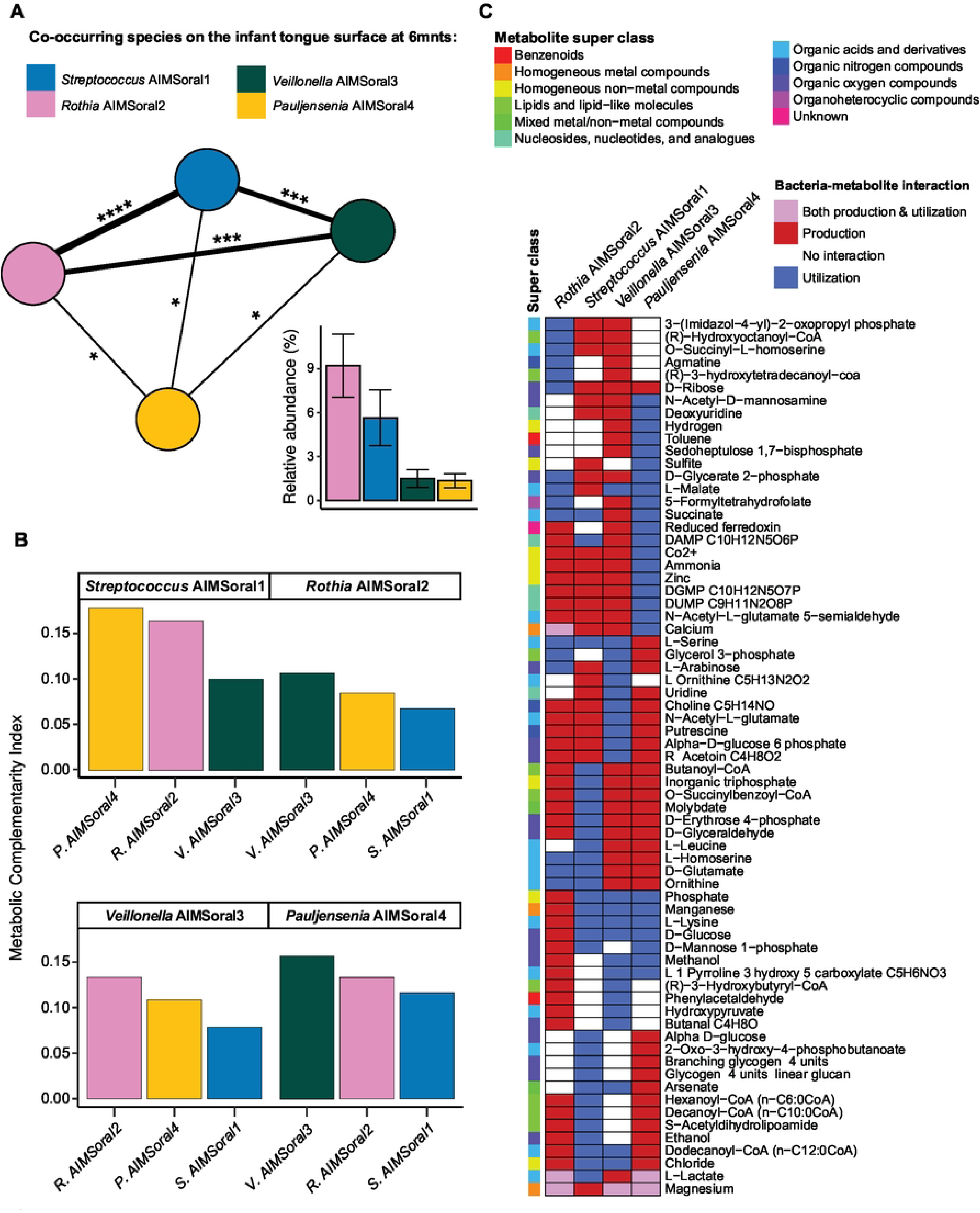
Metabolic complementarity and interactions within an infant tongue microbial co-occurrence network. **A)** Microbial network of four species (*Streptococcus* AIMSoral1=blue, *Rothia* AIMSoral2=pink, *Veillonella* AIMSoral3=green, *Pauljensenia* sp9002372285=yellow), sharing significant abundance correlations among each other on the tongue surface of six-month-old infants, and their relative abundances in this niche at this age. Significant correlations were calculated using Spearman’s rank correlation. Edges width indicate the strength of the significance. \**p*≤0.05, *p*≤0.01,\*\*\**p*≤0.001, \*\*\*\**p*≤0.0001. **B)** Bargraphs show the metabolic complementarity index for pairwise interactions among species in the tongue microbial network. Each facet title indicates the reference species for which the index is calculated. **C)** Heatmap showing bacterial-metabolite bipartite interactions (production=red, utilization=blue, both=purple, no interaction=white) potentially occurring among species in the tongue network. Metabolites are classified by their superclasses.

GEMS predicted distinct, species-level patterns of metabolite production and utilization, which are summarized in Fig. 5C and Table S14. Within the co-occurrence network, *Streptococcus* AIMSoral1 and *Rothia* AIMSoral2 were predicted to interact through 27 metabolites (Fig. 5C). Nine were potentially produced by *Streptococcus* AIMSoral1 and required by *Rothia* AIMSoral2, including malate. Malate exchange was supported by malate efflux carriers in *Streptococcus* AIMSoral1 (BLAST protein identity=95.2±1.6%; Table S11) and malate symporters in *Rothia* AIMSoral2 (51.6±0.2%; Table S12). The remaining 18 metabolites, including ethanol and lysine, were potentially produced by *Rothia* AIMSoral2 and required by *Streptococcus* AIMSoral1. Lysine exchange was supported by lysine exporters (LysE, K06895, Table S8) and amino acid permeases (98.8±1.5%; Table S11) in all *Rothia* AIMSoral2 and 93% of *Streptococcus* AIMSoral1 genomes, respectively. Although no metabolic interactions were exclusive to *Streptococcus* AIMSoral1 and *Rothia* AIMSoral2, both species exhibited unique interactions with other members of the network. Notably, *Veillonella* AIMSoral3 produced agmatine, utilizable only by *Rothia* AIMSoral2, while *Streptococcus* AIMSoral1 could use 2-oxo-3-hydroxy-4-phosphobutanoate produced by *Pauljensenia* AIMSoral4.

## DISCUSSION

In this study, we performed shotgun metagenomic sequencing on oral samples from 24 mother-infant pairs from the prospective AIMS cohort, enabling finer taxonomic resolutions than previous studies. This approach allowed the identification of previously undescribed species in the developing oral ecosystem and reconstruction of their genomes, revealing functional potential and metabolic interactions within the infant oral microbiome.

Our findings highlight the early dominance of *Streptococcus* spp. and increasing bacterial richness over time, including the emergence of the *Rothia* genus. Species-level changes in these genera, resulted in the establishment of *Streptococcus* AIMSoral1 and *Rothia* AIMSoral2 as predominant members of the infant oral microbiome. These genus-level transitions likely reflect adaptations to dietary changes during weaning and structural changes from dentition (40,41) aligning with previous 16S rRNA gene amplicon sequencing studies (6,8,40,41). However, our species-level observations diverge from previous findings, which suggested *R. mucilaginosa* and *S. salivarius* were highly prevalent after first dentition (6). Conversely, six-months-old AIMS infants lacked previously reported *S. salivarius* group species and had comparatively low prevalence of *R. mucilaginosa*. This discrepancy likely stems from lower 16S rRNA resolution, potentially misannotating *Streptococcus* AIMSoral1 and *Rothia* AIMSoral2 as *S. salivarius* and *R. mucilaginosa*, respectively, while our analysis shows these as novel species. We found no significant evidence that milk or formula feeding influenced the abundance of these species. To understand the ecological significance of these taxonomic shifts, we examined the functional capabilities of these predominant species, beginning with *Streptococcus* AIMSoral1.

*Streptococcus* AIMSoral1 displayed an enriched repertoire of genes for amino acid metabolism, particularly arginine and glutamate derivatives biosynthesis, suggesting adaptations to milk-derived nutrients as well as a role in oral nitrogen cycling (42–44). Additionally, dextran-degrading enzymes (GH66) may confer competitive advantages by breaking down EPS from other *Streptococcus* spp. (45,46), potentially facilitating colonization of mucosal and dental surfaces. The presence of the fimA adhesin suggests an increased specificity for the oral cavity (47). Notably, this gene was absent in *S. salivarius* and other infant-associated streptococci, except *S. lactarius*, the predominant streptococcal species in six-month-old infants’ oral cavities.

Our genomic analyses of *Rothia* AIMSoral2 suggest a metabolic strategy focused on rapid membrane synthesis (48) through enrichment of genes involved in fatty acid biosynthesis initiation compared to other infant-associated *Rothia* species. This, may in turn support environmental adaptation within the developing oral cavity (49). Genes encoding for GT11 cazymes (α(1,2/3)-fucosyltransferases) were also found to be enriched across *Rothia* AIMSoral2 genomes, potentially playing a role in host surface attachment (50,51). Together, these genomic features suggest *Streptococcus* AIMSoral1 and *Rothia* AIMSoral2 are adapted to the developing oral cavity, with distinct metabolic strategies providing ecological advantages during early colonization.

The strong positive correlation between *Streptococcus* AIMSoral1 and *Rothia* AIMSoral2 at six months suggests a potential cooperative relationship. Genomic and functional analyses suggest *Streptococcus* AIMSoral1 to create niches supporting *Rothia* AIMSoral2 colonization, modifying dextran-rich biofilm matrices. Meanwhile, *Rothia* AIMSoral2, encodes laccase-like multicopper oxidases (AA1_1/2), which can neutralize reactive oxygen species (ROS) (52). This capability may confer resilience against oxidative stress generated by hydrogen peroxide (H₂O₂) produced by cohabiting *Streptococcus* species (53) during biofilm formation (54,55). The presence of CBM50 domains further suggests that *Rothia* AIMSoral2 can effectively interact with other bacterial cell walls (56), consistent with Rothia’s role as an initiator of cell-cell interactions in early oral biofilms (57).

Metabolic modeling supports a cooperative relationship between *Streptococcus* AIMSoral1 and *Rothia* AIMSoral2. In our bacterial co-occurrence network analysis, *Streptococcus* AIMSoral1 exhibited the comparatively lowest metabolic complementarity with the other members, but highest complementarity with *Rothia* AIMSoral2 and *Pauljensenia* AIMSoral4. While previous studies have shown beneficial interactions between these genera (17,58,59), interactions between these undescribed species in the infant oral cavity remained uninvestigated. Metabolic modeling identified 27 potential interactions between *Streptococcus* AIMSoral1 and *Rothia* AIMSoral2, including malate and lysine, supported by genes encoding enzymes for their transport. These metabolic exchanges may underpin their stable co-occurrence in the infant oral microbiome, serving as energy source (60) and supporting essential cellular processes (61).

The identification of novel, prominent *Streptococcus* and *Rothia* species with specialized functional traits has significant implications for understanding early oral microbiome development and health outcomes. Arginine production by *Streptococcus* AIMSoral1 may modulate oral pH, as arginine catabolism by other bacteria generates ammonia (62,63), potentially neutralizing acids from cariogenic bacteria and reducing caries risk (64,65). Similarly, *Rothia* AIMSoral2 and *Veillonella* AIMSoral3 were predicted to produce ammonia, suggesting a broader microbial strategy for counteracting acidification and maintaining a balanced oral environment (5,66). Additionally, the exchange of lysine and agmatine within the network could influence pH regulation through polyamine (e.g. putrescine) metabolism, reinforcing metabolic interdependencies among these species. Beyond oral health, nitrate metabolism by *Rothia* species is linked to NO production through the enterosalivary pathway, a process benefiting systemic cardiovascular health (5,23). Disruptions in ammonia-producing pathways, due to antibiotics or diet, could therefore increase risk of caries and other oral diseases.

Genome-resolved metagenomics is a powerful tool for identifying the functional elements of the microbiome and provide insights into potential ecological interactions of *Streptococcus* and *Rothia* species during early oral microbiome development. However, MAGs represent incomplete genomes, potentially misrepresenting gene content. Most *Streptococcus* AIMSoral1 MAGs had medium-quality, which might have hindered functional annotations and downstream interpretations. Nevertheless, these MAGs clustered consistently with high-quality reference genomes in both gene presence/absence patterns and phylogenomic relationships, supporting our observations. Despite limitations, MAG reconstruction and metabolic modelling revealed the functional potential of undescribed oral species, generating testable hypotheses about species interactions for future infant oral microbiome studies.

Identifying novel species and their metabolic interactions has implications for understanding early-life oral microbiome assembly and health. Future research should prioritize the isolation of *Streptococcus* AIMSoral1 and *Rothia* AIMSoral2 and validate their predicted interactions, in particular regarding their pH homeostasis capabilities. Additionally, longitudinal studies that track these species from infancy through childhood will help clarify their long-term health impacts while revealing factors that influence their abundance dynamics. By deepening our knowledge of these ecological interactions, we ultimately can facilitate the development of precision-based interventions that promote both oral and systemic health.

## Material and methods

### Amsterdam Infant Microbiome Study (AIMS) cohort

AIMS (https://aimsonderzoek.nl/en/) (67) is a microbiome-focused multi-ethnic, prospective birth cohort study by the GGD Amsterdam (Public Health Service of Amsterdam, the Netherlands) within the Sarphati Cohort (68). The study was approved by the Medical Ethical Examination Committee of Amsterdam University Medical Center (METC Amsterdam UMC, Reference NL64399.018.17), with written informed consent obtained from all participants, and for infants, parental consent was secured after birth.

### Study population and sample collection

We collected 83 non-invasive oral biosamples (tongue swab: n=45, dental plaque: n=38) for metagenomic shotgun sequencing from 24 AIMS mother-infant pairs. Mothers self-collected both their samples (at 34 weeks pregnancy) and their infants’ (at 1 and 6 months postpartum). Tongue samples were collected by stroking a synthetic swab (FLOQSwabs®, Copan Italia s.p.a.) across the tongue dorsum for 30 seconds. Dental plaque was collected by using a sterile toothbrush (without toothpaste) to brush the buccal, lingual and occlusal surfaces of right lower molars for mothers and incisors in infants. Samples were home-stored in Liquid Amies Medium (eSwab®) at -18/-20°C, transported at -4°C, and stored at -70°C. Questionnaires tested in a face validity study (69) (Table S1) gathered metadata on infant milk-feeding type, antibiotic use, health status, and lifestyle factors (Table S1).

### DNA extraction and sequencing

DNA was extracted using the ZY-D4306 ZymoBIOMICS-96 Magbead DNA Kit (with the use of the KingFisher™ Flex Purification System for high throughput) following BaseClear protocols. Of 73 samples, 57 yielded sufficient DNA for sequencing: maternal (tongue=23, dental plaque=21), 1-month infants (tongue=9, dental plaque=0), and 6-month infants (tongue=18, dental plaque=14). Metagenomic libraries were sequenced on an Illumina NovaSeq platform, yielding 2x150bp paired-end reads.

### Pre-processing of metagenomic reads

Raw metagenomics reads were pre-processed using BBDuk and BBMap (70). Raw reads were trimmed to remove adapters, poor quality bases (Q15) and reads shorter than 45bp (minlength=45). BBMap was used to discard human reads (minID=0.95) and microbial reads were merged into longer single reads using BBMerge (min overlap=16). The SPAdes (71) assembler (metagenomic mode) was used for metagenomic assembly.

### MAG reconstruction

We used SemiBin for multi-sample binning to generate MAGs (SemiBin 1.0.0) (72). For each mother-infant pair, we mapped short reads to a set of assembled contigs (500bp) from the same mother-infant pair via Bowtie2 (73). Samples were binned in multi-sample binning mode (*SemiBin multi_easy_bin*).

MAG quality was estimated using CheckM (73,74) and GUNC v.0.1 (75), and genomes were taxonomically classified using the GTDB database. High-quality *Streptococcus* and *Rothia* MAGs were further passed through the *anvi-estimate-genome-completeness* module in Anvi’o v.8 (76) to estimate their completeness and redundancy; draft genomes were considered suitable for downstream analyses when displaying ≥50% completeness & ≤10% redundancy (i.e medium, medium-high and high quality). The complete list of *Streptococcus* and *Rothia* MAGs used in this study are provided in Table S5.

### Public reference genome acquisition

We conducted comparative genomic analyses on MAGs and publicly available *Streptococcus* and *Rothia* genomes from Progenomes3 (77) and GTDB (78). To ascertain the taxonomic placement of the undescribed *Streptococcus* species, we retrieved reference sequences from the *salivarius* group (*S. salivarius*: n=6, *S. vestibularis*: n=5, *S. thermophilus*: n=5). For *Rothia*, we used type strains sequences or GTDB representatives (Table S5). To perform comparative genomic analyses, we included only *Streptococcus* and *Rothia* species considered infant-associated, defined as those with average relative abundance of ≥5% and a prevalence of ≥30% (Figure S2). After quality filtering, 33 *Streptococcus* and 63 *Rothia* MAGs and genomes (hereon referred to as genomes) were used for genetic relatedness analysis, while 57 *Streptococcus* and 53 *Rothia* genomes comprised the final dataset for phylogenomics, pangenomics and metabolic enrichment analyses.

### Metagenomic species-profiling and strain tracking

In order to assess the relative abundance of bacterial taxonomic groups per sample, concatenated reads were mapped onto reference genomes from the GTDB R220 database (79) by using the computational tool sylph v.0.6.1 (80).

Single nucleotide variant (SNV) calling was used to determine overall strain-tracking between mothers and 1mo infants (i.e. vertical transmission) and mothers and 6mo infants (i.e. persisted vertical transmission or horizontal transmission). Reference-based strain tracking was performed using *inStrain* v1.0.0 (81) with reference genomes from the UHGG database, which we used to recruit quality-controlled merged reads using Bowtie2. Scaffold-level microdiversity metrics were calculated using the *inStrain profile* module. Pairwise species comparisons were performed using the *inStrain compare* module to calculate genome-level differences between samples (average nucleotide identity threshold = 0.999).

### Phylogenomic and genetic relatedness analyses

Phylogenomic analyses were carried out in Anvi’o v8. Amino acid sequences of single-copy core genes (https://merenlab.org/2017/06/07/phylogenomics/) were extracted from the genus set of 110 and 63 *Streptococcus* and *Rothia* genomes (see Methods: Public reference genomes acquisition), respectively, and aligned using the *anvi-get-sequences-for-hmm-hits* program. We further retrieved and aligned single-copy core genes from 33 genomes from the *salivarius* group for a targeted phylogenomic analysis of these *Streptococcus* species. We visualized the phylogenomics tree by using the resulting concatenated amino acid alignment as input for the *anvi-gen-phylogenomics-tree* program.

Genetic relatedness analyses were conducted for *Rothia* and the *salivarius* lineage within *Streptococcus* genus using the *anvi-compute-genome-similarity* module, which uses FastANI to calculate the average nucleotide identity (ANI). Genomes with an ANI ≥95% were classified as the same species.

### Streptococcus and Rothia pangenome

Pangenome analyses were carried out in Anvi’o v8. Genomes were converted into contig databases (*anvi-gen-contig-database*) and functionally annotated from KEGG databases (*anvi-hmm* and *anvi-run-kegg-kofams*). Functionally annotated contig databases were stored (*anvi-gen-genomes-storage*) and used for pangenome reconstruction. Using the *anvi-pangenome* and *anvi-interactive* modules we reconstructed the *Streptococcus* and *Rothia* genus pangenome and subsequently visualize the hierarchical clustering of genomes based on gene presence/absence (Method: Ward; Distance: Euclidean; Fig. S7 and Fig. S8).

### Metabolic enrichment analyses

We analyzed high- and medium-quality genomes to identify metabolic differences among *Streptococcus* and *Rothia* species. Functional annotations were used to predict metabolic pathways (KEGG MODULE resource) for *Streptococcus* and *Rothia* species. For this purpose, we used the *anvi-estimate-metabolism* module, which predicts both presence and completeness of the detected metabolic modules . Module abundance was determined as detailed in Watson et al. 2023 and the anvi’o tutorial https://anvio.org/m/anvi-estimate-metabolism (82). Enrichment of metabolic modules across species was estimated using the *anvi-compute-metabolic-enrichment* module, resulting in enrichment scores and adjusted p-values (q-value) of modules found in *Streptococcus* and *Rothia* genomes (83).

### Profiling of carbohydrate-active enzymes and adhesion potential

High-quality infant-type *Streptococcus* and *Rothia* genomes were annotated to the Carbohydrate-Active enZYmes (CAZy) database using Anvi’o v8 (run_dbCAN2) (84). For adhesion potential, CAZy annotation identified carbohydrate-binding modules (CBMs), while lectin-encoding genes were determined using known *Streptococcus* adhesins from literature and LectomeXplore (85). BLASTp (sequence identity≥40%, bitscore≥50, e-value≤0.001) (86) aligned genomes against lectin-sequences. Both CAZy and adhesin profiles were analyzed in R using hierarchical clustering (euclidean distance, Ward.D2 method) on scaled absolute gene counts. Differences in gene abundance across clusters were tested using Kruskal-Wallis followed by pairwise Wilcoxon Signed-Rank tests (*pairwise.wilcox.test*) for post-hoc comparisons.

### Metabolic complementarity/interaction potential

To test whether species co-occurrence was associated with metabolic interactions, we reconstructed phylogenetically-adjusted GEMS using PhyloMInt (v0.1.0) (87). Briefly, PhyloMInt employs CarveMe (26) to reconstruct GEMS and calculates (asymmetric) metabolic complementarity indices (MI_complementarity_) for genomes pairs based on the metabolites required or produced by their metabolic modules. We calculated the average MI_complementarity_ for all species pairs tested differences using pairwise-wilcoxon tests. To predict metabolic exchanges, we used the PhyloMInt PTM tool in iNAP 2.0 (88).

To support predicted metabolic interactions, we determined the presence of specific transporters (Table S9) for metabolites of interest using BLASTp (v2.6.0+) filtered by bitscore (≥50), e-value (≤0.001), sequence identity (≥40%) and best hit.

### Data analysis and statistics

Data analyses and statistical procedures were performed using a combination of RStudio (v2022.02.0+443) and Anvi’o (v8). Pangenomic hierarchical clustering, metabolic enrichment analyses using anvi-modules were performed in the Anvi’o 8.0 platformwhile other statistical tests were conducted in RStudio. False discovery rate (‘fdr’) correction was applied for multiple testing.

## Acknowledgements

We thank all the families who participate in the Amsterdam Infant Microbiome Study (AIMS) cohort and the entire research team responsible for the recruitment of the cohort and collection of samples and questionnaires.

## Data availability

The raw-reads for each AIMS biosample were submitted to the European Nucleotide Archive (ENA) at EMBL-EBI repository and the National Center for Biotechnology and Information (NCBI with BioProject accession number PRJEB88622). The dataset will be publicly available on 2025-09-01.

The code generating the main figures and underlying data are available at GitHub: https://github.com/nicholaspucci/infant-oral-microbiome-strep-rothia.

---

**Figure S1. Overview of the number of metagenomics reads retained per sample.** For each bar, colors indicate the proportion of reads fromA) tongue and B) tooth biofilm samples that passed (green) and failed (red) quality control as well as reads mapping to human DNA (blue). Failed and human reads were subsequently filtered out the dataset.

**Figure S2. Abundance and prevalence thresholds used for inclusion in the Spearman’s rank correlations and to define infant-associated Streptococcus species in our study.** Spearman’s rank correlation analyses were performed separately on bacterial species pairs on the tongue dorsum of A) 1-and B) 6-month-old infants and the C) dental plaque of 6-month-old infants that had a mean relative abundance of ≥1% (black dotted line). For the comparative genomics analyses, we defined infant-associated Streptococcus species as those showing a mean relative abundance ≥5% (horizontal red dotted line) and prevalence across AIMS infants ≥30% (vertical red dotted line) on the tongue dorsum of A) 1-and/or B) 6-month-old infants and/or in the C) dental plaque of 6-month-old infants.

**Figure S3. Oral microbiota of AIMS infants and corresponding milk feeding information. A)** PCoA of tongue dorsum (circles) and dental plaque (triangles) microbiomes from AIMS infants (one month=small; six months=large). For each sample, the corresponding milk feeding type is reported (breastmilk=pink, mixed=red,formula=green). PERMANOVA analyses showed no significant clustering based on milk feeding within the six months samples (adonis: R^2^=0.07, F=1.10, p=0.32). **B)** Relative abundances of Streptococcus AIMSoral1, Rothia AIMSoral2, Veillonella AIMSoral3 and Pauljensenia AIMSoral4 found in infants breast-, formula- and mixed milk-feeding. Significance among groups was tested for each species by pairwise Wilcoxon’s signed rank tests. No significant differences were detected. **C)** Correlation plot of Streptococcus AIMSoral1 and Rothia AIMSoral2 showed no association between their abundance and milk-feeding type.

**Figure S4. Heatmaps showing Spearman’s correlation coefficients for infant oral species as 1 and 6 months of age.** Correlations between oral species found to be abundant (mean relative abundance ≥1%) in the tongue biofilms of A) 1 and B) 6-months-old and C) dental biofilm of six-month-old AIMS infants are colored in red (positive) or blue (negative). Significance is indicated by asterisks: *p≤0.05, **≤0.01, ***≤0.001, ****≤0.0001. Self-self correlations are shown in grey.

**Figure S5. Phylogenomic tree of the *Streptococcus* genus.** Phylogenomic analysis of 110 *Streptococcus* genomes representing 89 species, including 89 reference genomes and 21 metagenome-assembled genomes (MAGs). *Lactococcus lactis* was included as an outgroup. Red circles indicate bootstrap values≥0.9.

**Figure S6. Heatmaps of Average Nucleotide Identity (ANI) of Streptococcus (salivarius group) and Rothia genomes.** Pairwise ANI comparisons between A) Streptococcus (salivarius group) and B) Rothia MAGs (white) and reference genomes (red). For each genome, the GTDB/AIMSoral and final species assignments are provided.

**Figure S7. Pangenome of infant-associated Streptococcus species in the oral cavity of AIMS infants.** The figure shows the pangenomic characteristics of infant-associated Streptoccoccus species sorted by gene presence/absence. The pangenome includes MAGs reconstructed from AIMS oral samples (sampling location: tongue dorsum = purple, tooth plaque = orange) collected at different timepoints (infant 1 month=green, infant 6 months = red). No high- and medium-quality MAGs from infant-associated species were retrieved from maternal oral samples. Reference genomes are given in grey. Species annotations as well as bars showing genome/MAG total length, completion and redundancy are shown for each strain. For each gene cluster, metrics such as number of genomes where a gene cluster is present (’num of contributing genomes) and various functional annotations are shown.

**Figure S8. Pangenome of infant-associated Rothia species in the oral cavity of AIMS infants.** The figure shows the pangenomic characteristics of infant-associated Rothia species sorted by gene presence/absence. The pangenome includes MAGs reconstructed from AIMS oral samples (sampling location: tongue dorsum=purple, tooth plaque=orange) collected at different timepoints (mother 34wks gestation = brown, infant 1 month =green, infant 6 months = red). Reference genomes are given in grey. Species annotations as well as bars showing genome/MAG total length, completion and redundancy are shown for each strain. For each gene cluster, metrics such as number of genomes where a gene cluster is present (’num of contributing genomes) and various functional annotations are shown.

**Figure S9. Metabolic complementarity and interactions among infant-associated Streptococcus and Rothia species in our study.** A) Bargraphs show the average metabolic complementarity index (±SD) for pairwise interactions among infant-associated Streptococcus and Rothia, and vice versa. For each complementarity species pair, the reference species is shown on the x-axis, while the other species is shown in the facet on top of the bar. B) Heatmap showing bacterial-metabolite bipartite interactions production = red, utilization = blue, both = purple, no interaction = white) potentially occurring among Streptococcus AIMSoral1, Rothia AIMSoral2 and their closely related infant-associated species in our study. Metabolites are classified by their superclasses.

**Figure S10.** Principal coordinate analysis (PCoA) of tongue dorsum microbiomes from AIMS mothers (34wks gestation=brown) and their infants (one month=green; six months=red).

**Figure S11. Analysis of 165 rRNA sequences from S. salivarius and Streptococcus AIMSoral1. Figure S12. Abundance of Streptococcus AIMSoral1 and Rothia AIMSoral2 across infant oral samples from Ferretti et al. 2018.**

**Table S1. Metadata for metagenomics samples retrieved from AIMS mothers (34wks gestation) and their infants (one and six months) available for this study.**

**Table S2. Overview strain tracking output (inStrain).**

**Table S3. Spearman rank-order R-values from pairwise abundance correlations between abundant (≥1%) species in the tongue dorsum and dental plaque samples of AIMS infants (one and six months).**

Table S4. Spearman rank-order *p*-values from pairwise abundance correlations between abundant (≥1%) species in the tongue dorsum and dental plaque samples of AIMS infants (one and six months).

**Table S5. Overview of medium- and high-quality metagenome-assembled genomes (MAGs) and genomes used for this study.**

**Table S6. Output of the metabolic enrichment analysis performed among infant-associated *Streptococcus* species.**

**Table S7. Output of the metabolic enrichment analysis performed among infant-associated *Rothia* species.**

**Table S8. Matrices of KEGG Orthologs (KOs) copy numbers per infant-associated *Streptococcus* and *Rothia* genomes/MAGs.**

**Table S9. Overview reference proteins used for genomic analyses of infant-associated *Streptococcus* spp. adhesin repertoire and validation of malate and lysine transporters from *Streptococcus AIMSoral*1 and *Rothia AIMSoral*2.**

**Table S10. Overview pairwise Wilcoxon test outputs on differences in gene copy-number among infant-associated *Streptococcus* and *Rothia* genomes/MAGs.**

**Table S11. NCBI BLASTp summary for malate and lysine transporter/permease proteins across infant-associated *Streptococcus* genomes/MAGs.**

**Table S12. NCBI BLASTp summary for malate and lysine uptake proteins across infant-associated *Rothia* genomes/MAGs.**

**Table S13. PhyloMInt complementarity indices for pairwise comparisons between infant-associated *Streptococcus* and *Rothia* genomes and *Veillonella* AIMSoral3 and *Pauljensenia AIMSoral*4.**

**Table S14. Overview metabolites predicted to be produced/utilized by *S.* salivarius, *Streptococcus* AIMSoral1, *R.* mucilaginosa, *R.* mucilaginosa_A, *Rothia* AIMSoral2, *Veillonella* AIMSoral3 and *Pauljensenia* AIMSoral4.**

**Table S15. AIMS oral microbiome data.**

